# Insights into the complex formation of a trimeric autotransporter adhesin with a peptidoglycan-binding periplasmic protein

**DOI:** 10.1101/2023.12.21.572085

**Authors:** Shogo Yoshimoto, Jun Sasahara, Atsuo Suzuki, Junichi Kanie, Kotaro Koiwai, Andrei N. Lupas, Katsutoshi Hori

## Abstract

Trimeric autotransporter adhesins (TAAs) are outer membrane (OM) proteins that are widely distributed in gram-negative bacteria and are involved primarily in adhesion to biotic and abiotic surfaces, cell agglutination, and biofilm formation. TAAs consist of a passenger domain, which is secreted onto the cell surface, and a transmembrane domain, which forms a pore in the OM to secrete and anchor the passenger domain. Because the interactions between TAAs and chaperones or dedicated auxiliary proteins during secretion are short-lived, TAAs are thought to reside on the OM without forming complexes with other proteins after secretion. In this study, we aimed to clarify the interactions between an *Acinetobacter* TAA, AtaA, and a peptidoglycan (PG)-binding periplasmic protein, TpgA. Pull-down assays using recombinant proteins identified the interacting domains. X-ray crystallography at 2.6 Å resolution revealed an A3B3 heterohexameric complex structure composed of the N-terminal domain of TpgA and the transmembrane domain of AtaA. TpgA-N consists of two short α helices and three antiparallel β strands, yielding an ααβββ topology similar to BamE. However, the regions corresponding to BamE interfaces with BamA and BamD differ in TpgA-N. All-atom molecular dynamics simulations and mutational assays revealed that both electrostatic and hydrophobic interactions contribute to stable complex formation. Bioinformatic analyses indicate that the TAA-TpgA complex occurs in a wide range of species. These findings will contribute to a better understanding of TAAs and the cell envelope.

**Importance:** Gram-negative bacteria have specialized secretion systems (SSs) that translocate molecules from the cytoplasm to the extracellular space. Type V SSs have a simpler structure consisting of a functional passenger domain and a transmembrane domain involved in the secretion and anchoring of the passenger domain. Here, we provide the first direct evidence that a trimeric autotransporter adhesin (TAA) exported by a type Vc system forms a stable complex with a peptidoglycan–binding periplasmic protein. The 2.6 Å structure of the A3B3 heterohexamer, together with simulation and mutational data, reveals complementary electrostatic and hydrophobic contacts that stabilize flexible loops on the periplasmic face of the TAA transmembrane barrel. Conservation of the *taa*–*tpgA* gene cassette and of key interface residues across diverse genera suggests that this coupling is a common strategy for tuning TAA stability and indicates that the envelope architecture of many TAAs, including those related to pathogenicity, is more elaborate than previously appreciated.

## Introduction

Cells of gram-negative bacteria are enveloped by the inner membrane, peptidoglycan (PG), and outer membrane (OM) (1). To translocate molecules from the cytoplasm to the extracellular space across these barriers, gram-negative bacteria have specialized secretion systems (SSs) known as type I to VI and VIII to XI (2–5). Molecules secreted through these SSs are responsible for various functions, including cell adhesion, immune evasion, toxin activity, stress response, motility, the transfer of genetic information, and substrate degradation. Therefore, understanding the structure and function of these SSs is important not only in microbiology but also for medical and bioengineering applications.

Type V SSs, known as autotransporters (6), have simpler architectures than that of the other types of SSs. These proteins can be further divided into subclasses, type Va to Ve SSs, which share a structure consisting of a passenger domain secreted into the extracellular space and a transmembrane domain involved in the secretion of the passenger domain through the OM and the anchoring of the passenger domain to the cell surface (Fig. S1)(7). In the type Vb SS only, the passenger and transmembrane domains are produced as separate molecules, and the other type V SSs are translated as a single molecule that contains both domains in the same polypeptide chain. In the type Va SS, which secretes monomeric autotransporters, the polypeptides synthesized in the cytoplasm pass through the inner membrane via the Sec system, and periplasmic chaperones, such as SurA, Skp, DegP, DnaK, and FkpA, assist this periplasmic transit (8–11). With the assistance of the β-barrel assembly machinery (BAM) complex, the transmembrane domain inserts into the OM as a 12-stranded β-barrel, and the passenger domain of the same molecule is transported to the extracellular milieu through a hybrid pore composed of the BAM complex and the transmembrane domain with no need for external energy sources (7, 12–14). The overall secretion process of the type Vc SS, another subtype of type V SSs that secretes trimeric autotransporter adhesins (TAAs) (15), is thought to be similar to that of the type Va SS, except that a trimeric intermediate is formed before membrane insertion (16). The passenger domain of TAAs usually remains on the cell surface after secretion due to anchoring by the transmembrane domain. In addition to the general periplasmic chaperones and BAM complex, in the type Vc SS, genes that form an operon with *taa* in the genome have also been reported to participate in the biogenesis of TAAs: the *Burkholderia* heptosyltransferase BimC participates in the localization of BimA to the cell pole (17), and the *Salmonella* inner membrane lipoprotein SadB participates in the outer membrane insertion of SadA (18, 19). The interaction between TAAs and chaperones or dedicated auxiliary proteins is transient (18), and TAAs are thought to reside on the outer membrane through its transmembrane domain without forming complexes with other proteins after secretion.

The gram-negative bacterium *Acinetobacter* sp. Tol 5 shows extremely high adhesiveness to various material surfaces, including hydrophobic plastics, hydrophilic glass, and metals, through its TAA, AtaA (20). The polypeptide chains of AtaA, consisting of 3,630 amino acids, form a very large homotrimeric fibrous structure that is more than 1 MDa in molecular weight and 260 nm in length (21). We previously reported that the *tpgA* gene, which forms a single operon with *ataA*, is involved in the biogenesis of AtaA (22). TpgA is characterized as a periplasmic protein composed of the SmpA/OmlA-like N-terminal domain and the OmpA_C-like C-terminal domain containing a PG-binding motif. The interaction between TpgA and AtaA was shown by a pulldown assay without crosslinking (22), suggesting that TpgA forms a stable complex with AtaA rather than a transient interaction.

In this study, we aimed to clarify the interaction between TpgA and AtaA. This study provides structural insights into PG-binding periplasmic proteins that form stable complexes with TAAs.

## Results

### Identification of the interacting domains of AtaA and TpgA

To identify which region of AtaA binds to TpgA, we performed a pull-down assay using recombinant AtaA fragments containing several AtaA domains, as shown in Fig. 1A. AtaA_723-806_, AtaA_1221-1416_, AtaA_2215-2395_, and AtaA_2777-2903_ were designed to connect to both the N- and the C-terminal trimeric GCN4 adaptors, GCN4pII, which have been used to stabilize coiled-coil domains of TAA recombinant proteins (23). AtaA_3524-3630_ was designed to be fused to the N-terminal OmpA signal sequence for secretion and presentation on the outer membrane in *Escherichia coli* because it contains the transmembrane domain. We also used previously constructed expression plasmids for AtaA_59-325_, AtaA_2905-3168_, and AtaA_3170-3561_ (24, 25). All of these recombinant proteins were connected to a C-terminal His-tag except for AtaA_3524-3630_ which was connected to the N-terminal signal sequence followed by His-tag. Considering the repetitive sequences, these recombinant proteins cover almost the entire region of AtaA. The cell lysate of Tol 5 Δ*ataA*, which contained native TpgA, was mixed with recombinant AtaA fragment-bound nickel-nitrilotriacetic acid (Ni-NTA) beads, and the pulled-down proteins were subjected to SDS‒PAGE and detected by CBB staining and western blotting using an anti-TpgA antibody. All of the recombinant proteins of AtaA fragments immobilized on the beads were detected by CBB staining, but TpgA was detected only in the sample incubated with the beads to which AtaA_3524-3630_ had been bound (Fig. 1B). These results suggest that TpgA specifically binds to AtaA_3524-3630_, the C-terminal region containing the transmembrane domain.

**Figure 1.**
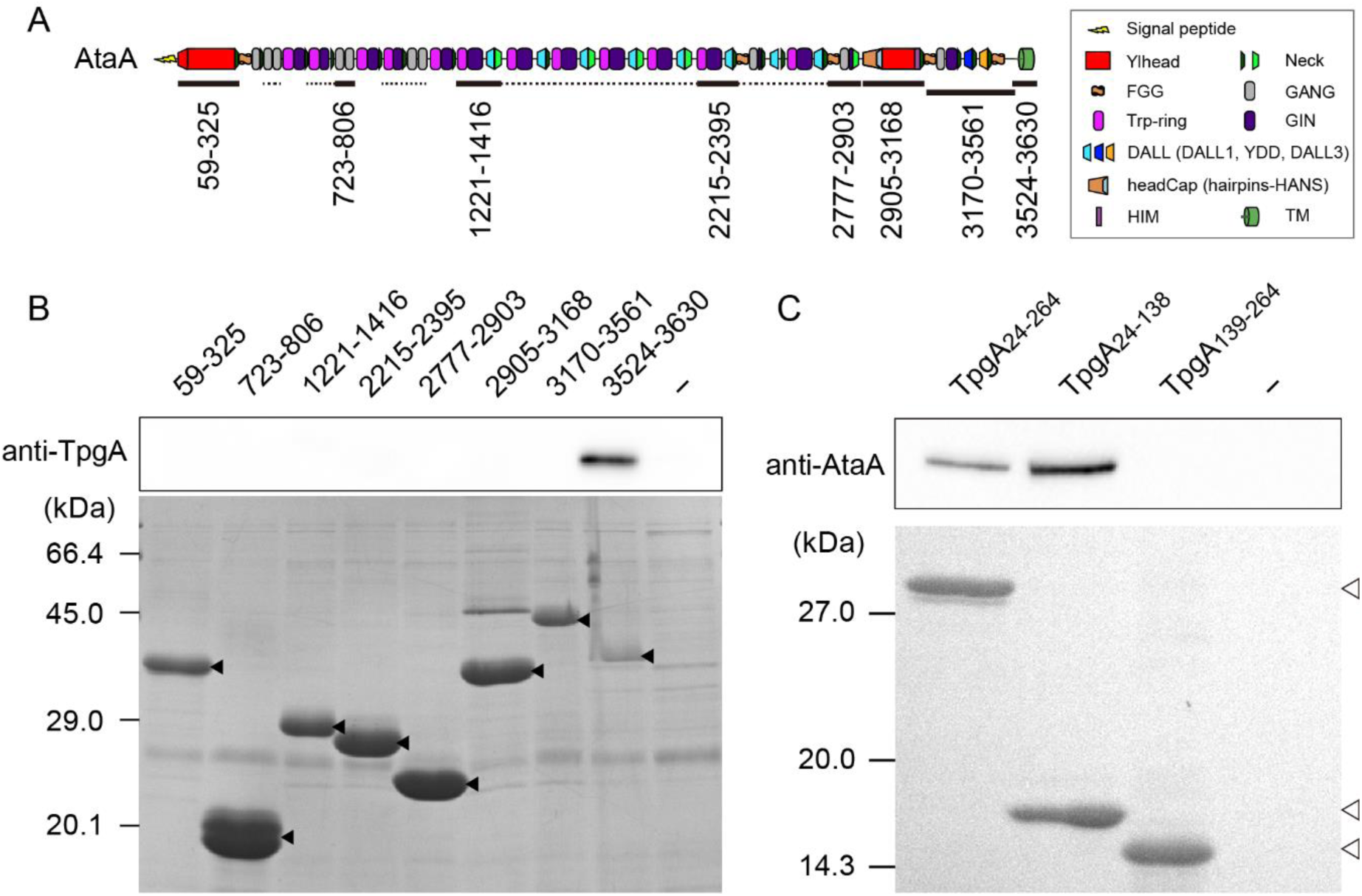
Identification of the interacting domains of AtaA and TpgA. (A) Schematic structure of AtaA. The black lines show the region where the recombinant protein was constructed, and the dotted lines show the region with high sequence similarity (>50%) to the recombinant proteins. (B) Pull-down assay using recombinant AtaA fragments and TpgA-containing cell lysate. The cell lysate of Tol 5 Δ*ataA* was mixed with recombinant AtaA fragment-bound Ni-NTA beads or untreated beads (-). The pulled-down proteins were subjected to SDS‒PAGE and analyzed by western blotting to detect TpgA and CBB staining to detect AtaA fragments. The solid triangles indicate recombinant AtaA fragments. The upper numbers indicate the amino acid numbers of each recombinant AtaA fragment immobilized onto Ni-NTA beads. (C) Pull-down assay using recombinant TpgA fragments and AtaA-containing cell lysate. The cell lysate of Tol 5 Δ*tpgA* was mixed with recombinant TpgA fragment-bound Ni-NTA beads or untreated beads (-). The pulled-down proteins were subjected to SDS‒PAGE and analyzed by western blotting to detect AtaA and CBB staining to detect TpgA fragments. The open triangles indicate recombinant TpgA fragments.

Next, to determine which domain of TpgA binds to AtaA, we performed a pull-down assay using recombinant TpgA or its domains. TpgA_24-264_ have been previously designed with its signal sequence removed. TpgA_24-138_ and TpgA_139-264_ consist of the N- and C-terminal domains of TpgA, respectively. All of these recombinant proteins were connected to a His-tag. The cell lysate of Tol 5 Δ*tpgA*, which contained native AtaA, was mixed with recombinant TpgA (or TpgA domain)-bound Ni-NTA beads, and the pulled-down proteins were subjected to SDS‒PAGE and detected by CBB staining and western blotting using an anti-AtaA antiserum. AtaA was pulled down with TpgA_24-264_ and TpgA_24-138_ but not with TpgA_139-264_ (Fig. 1C). These results suggest that the transmembrane domain of AtaA (AtaA- TM) forms a complex with the N-terminal domain of TpgA (TpgA-N).

### Crystal structure of the transmembrane domain of AtaA in complex with the N-terminal domain of TpgA

To determine the structure of AtaA-TM in complex with TpgA-N, we attempted to purify the protein complex of AtaA-TM and TpgA-N. The cell lysate of *E. coli* expressing His-tagged TpgA_24-138_ was added to a Ni-NTA column, which was washed to remove the unbound proteins. The OmpA signal sequence-fused AtaA_3524-3630_ without a His-tag was expressed in *E. coli,* and the membrane fraction solubilized by a detergent was added to the TpgA_24-138_-bound Ni-NTA column for the formation of the protein complex. After the unbound proteins were washed out, the protein complex was eluted with imidazole solution and further purified by size-exclusion chromatography. As shown in Fig. 2A and 2B, highly pure AtaA-TM in complex with TpgA-N was obtained at the elution peak corresponding to 82 kDa. This molecular weight was very close to that of three molecules of 11.1 kDa AtaA- TM and three molecules of 14.9 kDa TpgA-N (78 kDa), suggesting that AtaA-TM and TpgA-N form a stable A3B3 heterohexamer complex.

**Figure 2.**
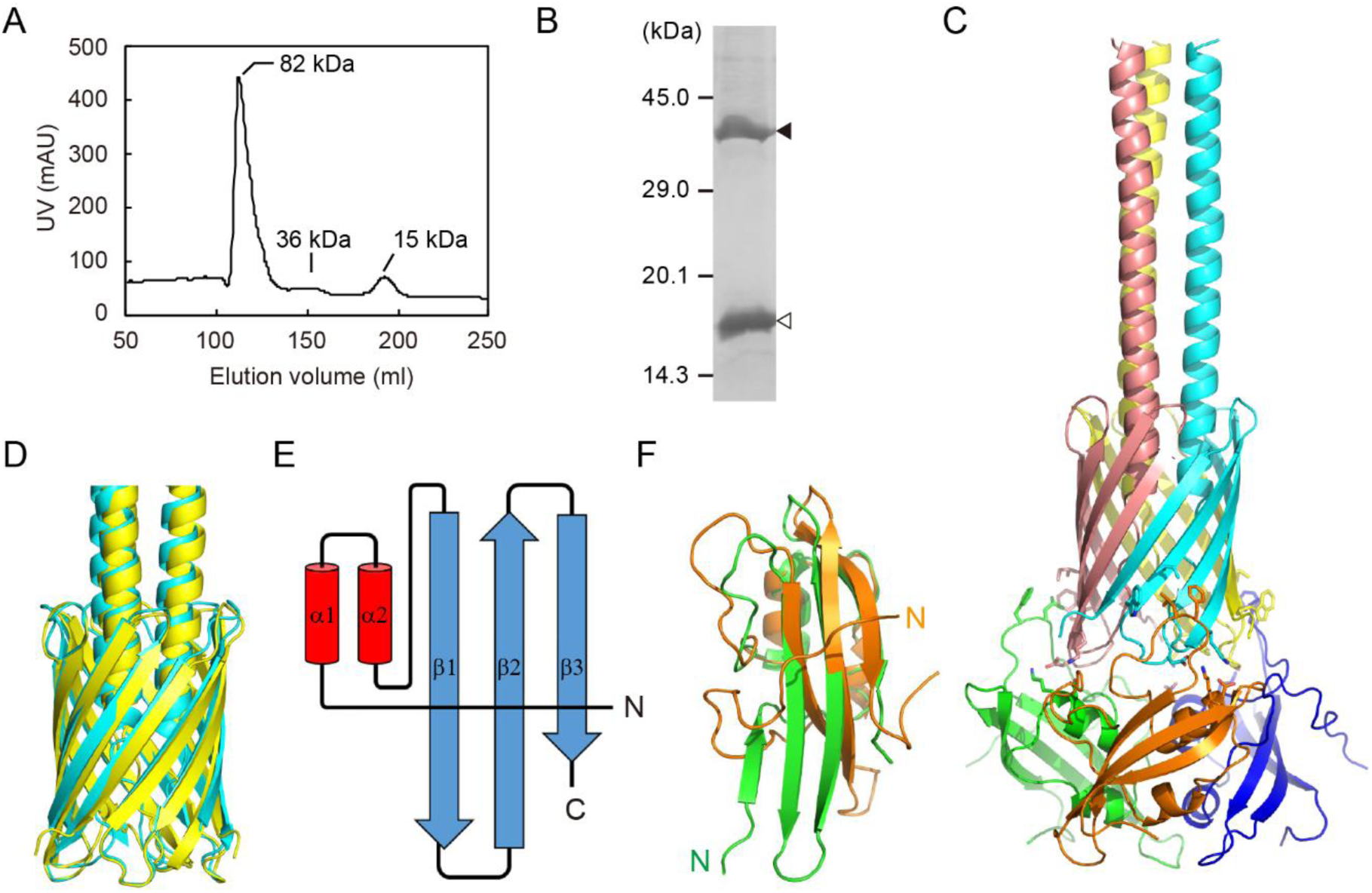
Crystal structure of the transmembrane domain of AtaA in complex with the N-terminal domain of TpgA. (A) Size-exclusion chromatography of the complex of AtaA- TM and TpgA-N. (B) The purified complex of AtaA-TM and TpgA-N was subjected to SDS‒PAGE followed by CBB staining. The solid and open triangles indicate the AtaA-TM trimer and TpgA-N, respectively. (C) Overall structure of the AtaA-TM and TpgA-N complex. The AtaA-TM monomers are shown in cyan, pink and yellow, and the TpgA-N monomers are shown in orange, green and blue, respectively. (D) Comparison of the AtaA-TM (yellow) with the transmembrane domain of *H. influenzae* Hia (2GR7; cyan). (E) A topology diagram of TpgA-N. α-helices are shown as cylinders, and β-strands are shown as arrows. (F) Comparison of TpgA-N (orange) with *E. coli* BamE (5D0Q; green).

After crystallization screens using the purified protein complex, we determined the structure of AtaA-TM in complex with TpgA-N at 2.6 Å resolution by X-ray crystallography (Fig. 2C, PDB ID: 9VNJ). This is the first report showing a quaternary structure of the transmembrane domain of TAA in complex with another protein. Three molecules of TpgA-N bind to the periplasmic side of the homotrimeric transmembrane β- barrel of AtaA-TM, forming the A3B3 heterohexamer, as estimated by size exclusion chromatography. The monomer of AtaA-TM contains an N-terminal helix followed by a four-stranded β-sheet, and a 12-stranded β-barrel is formed through homotrimerization. As shown in Fig. 2D, the structure of AtaA-TM is very similar to the previously determined structures of transmembrane domains of other TAAs, such as those of *Haemophilus influenzae* Hia and *Yersinia enterocolitica* YadA (26, 27). The structure of TpgA-N consists of two small α-helices and three antiparallel β-strands with an ααβββ topology (Fig. 2E).

The two α-helices pack against the β-sheet, forming an α/β 2-layer sandwich architecture. The most similar known protein is that of *E. coli* BamE (28). While they share 27% amino acid sequence identity, the structures differ with a root-mean-square deviation (RMSD) of 3.7 Å, particularly in the N-terminal region and in the β1- and β2-strands, which are involved in the binding of BamE to BamA and BamD (Figs. 2F and S2) (29).

### Interaction between the transmembrane domain of AtaA and the N-terminal domain of TpgA

To investigate how AtaA and its TpgA complex behave in the outer membrane, we conducted all-atom molecular dynamics (MD) simulations. The crystal structures of AtaA- TM and the complex of AtaA-TM and TpgA-N were embedded in an asymmetric lipid membrane: the upper leaflet consisted of lipopolysaccharides (LPS) derived from *Acinetobacter baumannii* and the lower leaflet composed of a mixture of 1-palmitoyl-2-oleoyl-*sn*-glycero-3-phosphoethanolamine (POPE) and 1-palmitoyl-2-oleoyl-*sn*-glycero-3- phosphatidylglycerol (POPG) in a 4:1 ratio, which mimicked the reported *Acinetobacter* lipid composition (30) (Fig. 3A). The structure of AtaA-TM reached equilibrium during the 500 ns simulation, and no significant difference was observed in its RMSD in the presence or absence of TpgA (Fig. S3). Calculation of per–residue RMSF of AtaA-TM showed that residues 3573 to 3576, which form the periplasm–exposed α–helix to β1 loop, and residues 3601 to 3604, which form the β2 to β3 loop, exhibited decreased RMSF upon TpgA binding (Fig. 3B). In YadA from *Y. enterocolitica*, the β2 to β3 loop has also been reported to be flexible (31). These data indicate that complex formation with TpgA stabilizes these loops of AtaA-TM. The loops are positioned adjacent to the hydrophobic tip of TpgA containing L80 and F81 and are in proximity to two electrostatic pairs, K64–D3601 and D65–K3576, which likely provide the stabilizing contacts (Fig. 3C). To examine the extent to which these interactions contribute to complex formation, we performed a pull-down assay using point-mutated TpgA-N proteins. The cell lysate of Tol 5 Δ*tpgA*, which contained native AtaA, was mixed with mutated TpgA-bound Ni-NTA beads, and the pulled-down proteins were subjected to SDS–PAGE and detected by CBB staining. As shown in Fig. 3D, the D65A and L80A mutations did not affect complex formation between AtaA-TM and TpgA-N, but the K64A and F81A mutations reduced complex formation. The K64A/D65A and L80A/F81A double mutations reduced the interaction. These results indicate that both electrostatic and hydrophobic interactions caused by these residues are important for complex formation.

**Figure 3.**
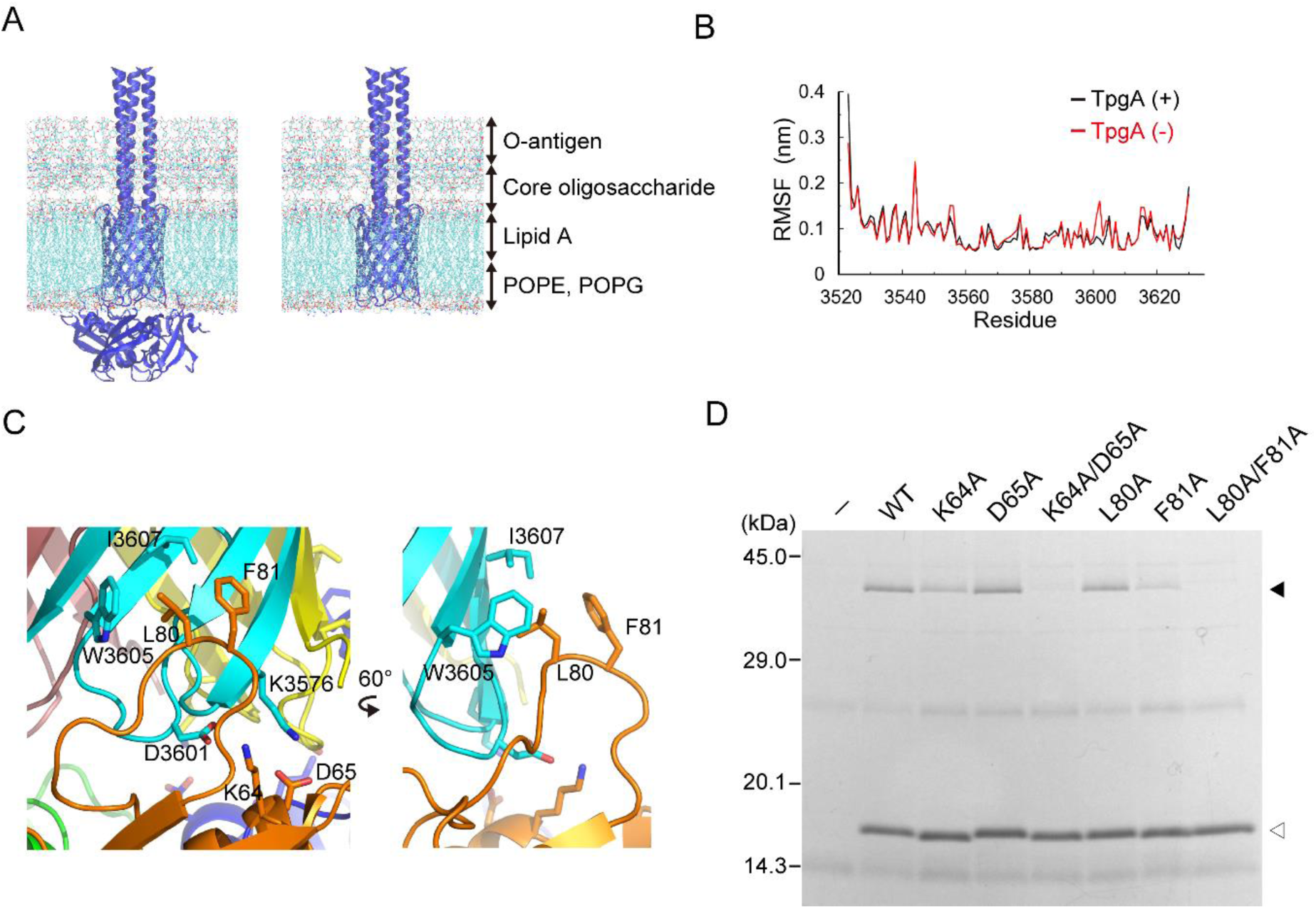
Interaction between the transmembrane domain of AtaA and the N-terminal domain of TpgA. (A) A snapshot from the MD simulation of AtaA-TM and TpgA-N. Polypeptides of AtaA-TM and TpgA-N are shown as cartoon models in blue, and phospholipids and lipopolysaccharides (LPS) are shown in line representation. (B) The root mean square fluctuation (RMSF) of each residue of AtaA-TM calculated from 500-ns simulation. (C) Enlarged views showing hydrophobic and electrostatic interactions between AtaA-TM (cyan) and TpgA-N (orange). The interacting residues are shown in stick representation. (D) Pull-down assay of AtaA-TM and mutated TpgA-N. Cell lysate containing AtaA-TM was mixed with His-tagged TpgA-N-bound Ni-NTA beads or untreated beads as a control (-). The pulled-down proteins were subjected to SDS‒PAGE and detected by CBB staining. The solid and open triangles indicate the AtaA-TM and point-mutated TpgA-N proteins, respectively.

### Bioinformatic analysis of the AtaA-TpgA interaction

To clarify whether the TAA-TpgA complex exists in a wide range of species, we performed bioinformatic analysis. The amino acid sequences with similarity to that of TpgA-N were collected using BLAST and clustered based on sequence similarity using CLANS (32). As shown in Fig. 4A, eight clusters were observed. TpgA-like sequences formed cluster I, which was different from clusters II and III containing a typical SmpA/OmlA family protein, BamE. Cluster I included sequences from a wide range of Pseudomonadota (Proteobacteria), such as *Altererythrobacter*, *Burkholderia*, *Neisseria*, *Acinetobacter*, *Stenotrophomonas*, *Escherichia*, and *Campylobacter*, as shown in Table S1.

**Figure 4.**
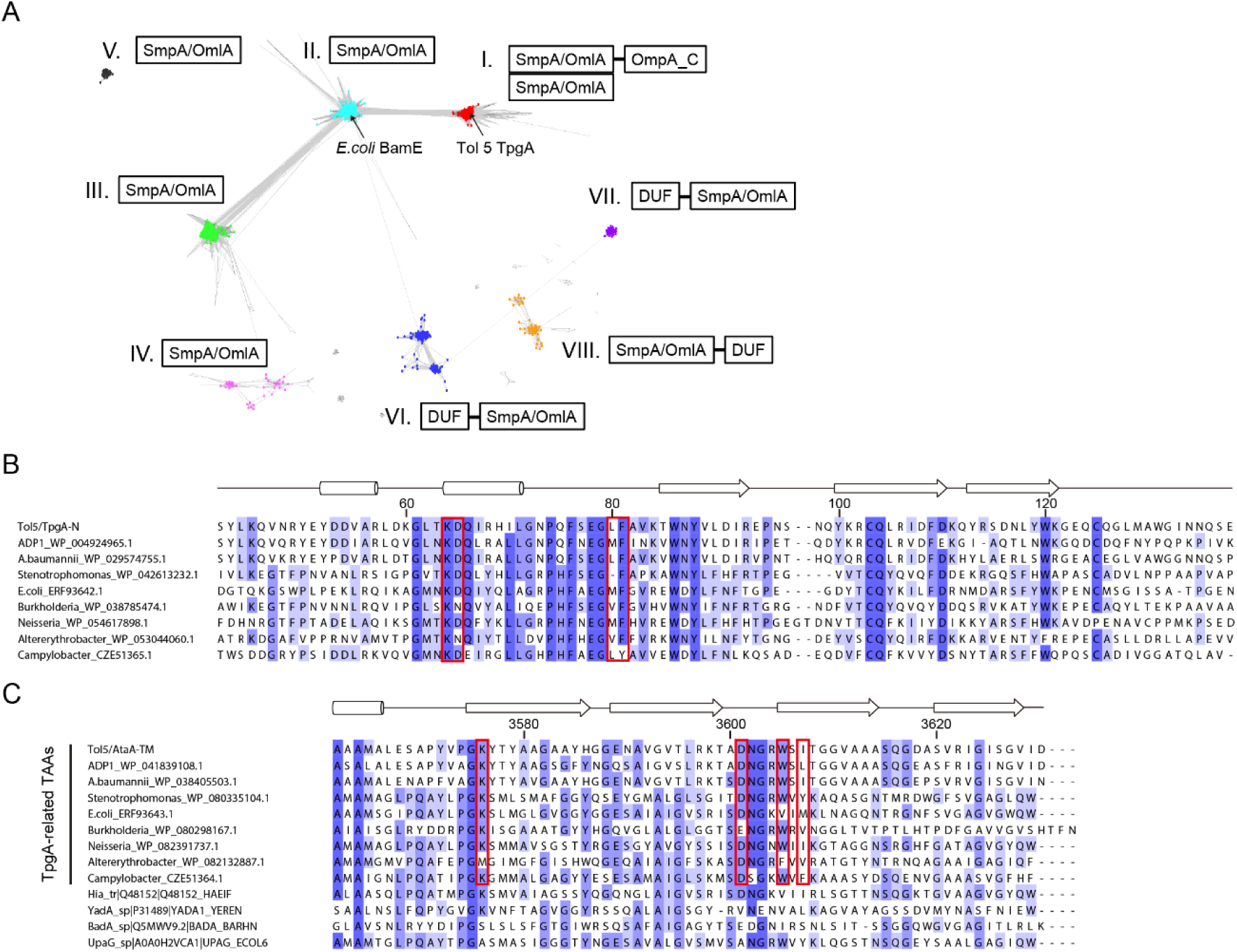
Bioinformatic analysis of the N-terminal domain of TpgA and the transmembrane domain of AtaA. (A) Cluster map of SmpA/OmlA family proteins. The map represents individual proteins and their sequence similarities as dots and gray lines, respectively. The positions of TpgA of Tol 5 and BamE of *E.coli* are indicated by arrows. (B) Amino acid sequence of the TpgA-N and TpgA-like proteins found in cluster I. Red boxes indicate residues that interact with AtaA. (C) Amino acid sequence of the C-terminal domain of AtaA; other TAAs found upstream of *tpgA*-like genes in the genome; and four representative TAAs (Hia, YadA, BadA, and UpaG) not related to TpgA. The red boxes indicate residues that interact with TpgA.

The domain composition of the sequences differed from cluster to cluster (Fig. 4A). In cluster I, approximately 80% of the sequences have an OmpA_C domain in the C-terminus, similar to that in TpgA, and most of the sequences in clusters II, III, IV, and V have no extra domain. Those in clusters VI and VII have a domain of unknown function (DUF) in the N-terminus, and those in cluster VIII have a DUF in the C-terminus. Thus, the domain structure of the sequences in cluster I, including TpgA, which consists of an N-terminal SmpA/OmlA domain and a C-terminal OmpA_C domain, is distinct and unique among proteins that contain the SmpA/OmlA domain.

In addition, 91% of the sequences in cluster I were downstream of the *taa* gene in the genome, although those in other clusters were rarely near the *taa* gene. This result is consistent with a previous finding that the occurrence of *ata*, a TAA of *Acinetobacter baumannii*, in the genome was tightly linked to that of *tpgA* (33). In the *taa-tpgA* gene cassette found in cluster I, L80 and F81 of TpgA were conserved as hydrophobic and aromatic residue pairs, respectively (Fig. 4B). Furthermore, K64 of TpgA and D3601 of AtaA, which interact with each other, are also conserved (Fig. 4B, C). In contrast, D3601 is not conserved in TAAs that do not form the *taa-tpgA* gene cassette, such as YadA (27), BadA (34), or UpaG (35), except for Hia (26), suggesting that D3601 of TAAs for which *tpgA* exists downstream in the genome, is conserved because of its interaction with TpgA. These new findings suggest that the TAA-TpgA protein complex is formed in a wide range of species.

## Discussion

TAAs interact with various auxiliary proteins during secretion but are thought to consist of simple components after secretion. In this study, we revealed that TpgA forms a heterohexamer with AtaA (Fig. 2). This is the first report directly showing that the transmembrane domain of TAA forms a stable complex with periplasmic proteins.

While each SS of type Va to Ve secretes a variety of passenger domains into the extracellular space or onto the cell surface, they share a common feature in that the transmembrane domain is anchored only to the OM, except for some type Ve SSs, which have a PG-binding LysM domain on the periplasmic side of their own polypeptide chain (Fig. S1) (36). AtaA does not have a PG-binding domain, but TpgA has a PG-binding domain containing the OmpA_C-like sequence at the C-terminus, and it was shown that TpgA actually isolated with PG (22). Considering that the N-terminal domain of TpgA binds to the C-terminal transmembrane domain of AtaA (Figs. 1, 2), these proteins likely form the AtaA-TpgA-PG complex (Fig. 5). In mammalian cells, the cell adhesion protein cadherin also forms a complex with actin filaments via catenins (37, 38). This implies that anchoring cell adhesion proteins to the structures involved in cell shape and strength, such as the cytoskeleton or peptidoglycan, would be universally important for mechanical stabilization.

**Figure 5.**
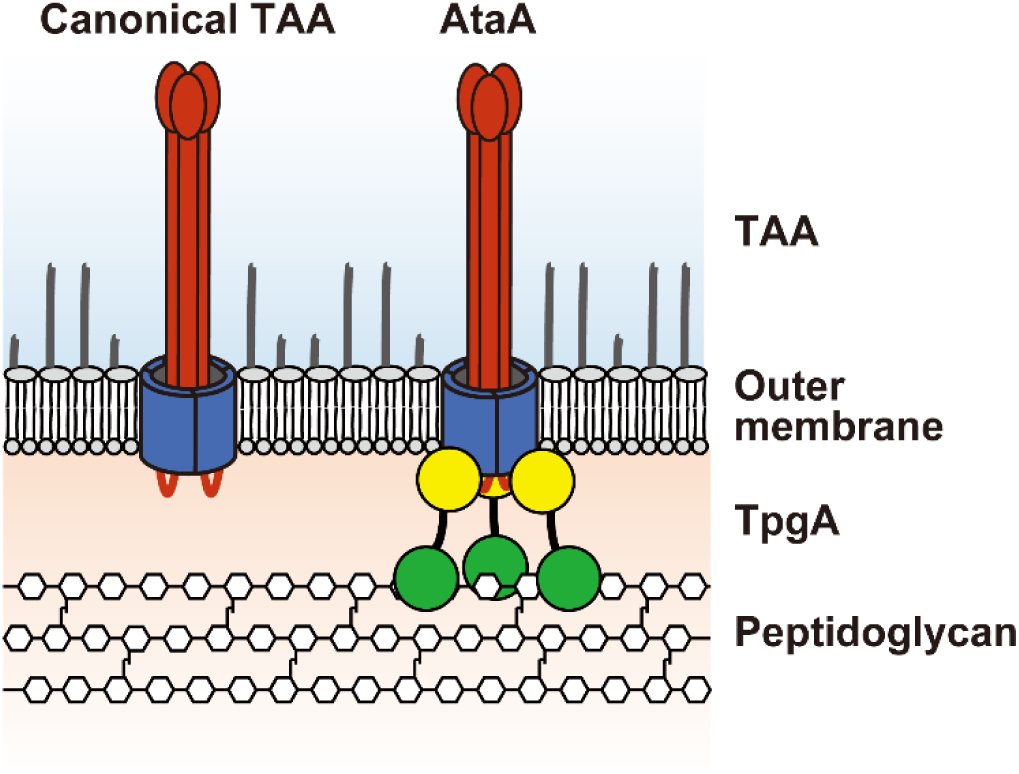
Schematic representation of a canonical TAA and the AtaA-TpgA- peptidoglycan complex. The passenger and transmembrane domains of TAAs are shown in red and blue, respectively. The N-terminal and C-terminal domains of TpgA are shown in yellow and green, respectively.

The overall structure of TpgA-N was similar to that of BamE, but the structures of the regions of BamE that bind to BamA and BamD (29) and the corresponding region of TpgA- N were different (Figs. 2F and S2). Furthermore, clustering analysis revealed that the *tpgA*-like genes formed a distinct cluster from other SmpA/OmlA family genes, such as *bamE* (Fig. 4A), and the TpgA residues interacting with AtaA were not conserved in BamE (Fig. S2). These results indicate that TpgA does not function as a subunit of the BAM complex. When TpgA was used as bait to isolate the protein complex of AtaA-TM and TpgA-N, the other β-barrel proteins were not co-isolated from *E. coli* lysate with TpgA-N (Fig. 2B). This indicates that TpgA does not bind nonspecifically to β-barrel proteins but rather specifically to AtaA, via the hydrophobic and electrostatic interactions (Fig. 3).

The *tpgA*-like genes were found widely in the strains belonging to Pseudomonadota (Proteobacteria), and most of them were located downstream of the *taa* genes in the genome (Fig. 4 and Table S1). These findings suggest that the TAA-TpgA protein complex is formed in a wide range of species. Notably, the clinically relevant pathogen *A. baumannii* also possesses a TAA-TpgA cassette as previously reported (33, 39), suggesting that TpgA may promote TAA-dependent virulence in pathogenic bacteria; dedicated experimental studies are still needed to confirm this possibility.

In conclusion, we revealed the structure of a PG-binding periplasmic protein that forms a stable complex with the transmembrane domain of a TAA. Our findings will contribute to a better understanding of the type Vc SS and the control of TAA-mediated adhesion, which is important for medical and engineering applications.

## Materials and Methods

### Bacterial strains and growth conditions

The bacterial strains and plasmids used in this study are listed in Table S2. *Acinetobacter* sp. Tol 5 and its derivative strains were grown in basal salt medium supplemented with toluene or in Luria–Bertani (LB) medium at 28°C, as described previously (20). *E. coli* DH5α, BL21(DE3), and their derivative strains were grown in LB medium supplemented with 100 μg/ml ampicillin for cells harboring pASK-IBA plasmids or with 50 μg/ml kanamycin for cells harboring pET plasmids at 37°C.

### Construction of plasmids

The primers used in this study are listed in Table S3. To construct plasmids expressing a recombinant AtaA fragment fused to a GCN4pII-tag at both the N- and C-termini and a His-tag at the C-terminus, DNA fragments encoding AtaA_723-806_, AtaA_1221-1416_, AtaA_2215-2395_, or AtaA_2777-2903_ were amplified via PCR and inserted into the BsaI site of the *E. coli* expression vector pIBA-GCN4tri-His (23). To construct a plasmid expressing AtaA_3524-3630_ fused to the His-tag at the N-terminus at the OM, the DNA fragment encoding this gene was amplified via PCR and inserted between the NheI and HindIII sites of the *E. coli* expression vector pASK-IBA12 (IBA Lifesciences, Göttingen, Germany), generating pIBA-His-AtaA_3524-3630_. To remove the His-tag sequence from pIBA-His-AtaA_3524-3630_, a DNA fragment was amplified via PCR using pIBA-His-AtaA_3524-3630_ as a template with two primers (TM-dHis-f/r) and then self-ligated, generating pIBA-AtaA_3524-3630_. To construct plasmids expressing TpgA_24-138_, TpgA_139-264_ or the point mutated TpgAs fused to a His-tag at the C-terminus, a DNA fragment was amplified via PCR using pET28d::TpgA_24-264_ (24) as a template with corresponding primer sets and then self-ligated, generating pET28d::TpgA_24-138_, pET28d::TpgA_139-264_, and plasmids for the point-mutated TpgAs.

### Protein purification

To purify the complex of TpgA-N and AtaA-TM, an overnight culture of BL21 (DE3; pET28d::TpgA_24-264_) was diluted 1:100 in LB medium and incubated for 3 h at 37°C. After incubation, isopropyl β-D-thiogalactopyranoside (IPTG) (final concentration: 0.1 mM) was added to the medium, and the culture was further incubated at 18°C for 18 h. The cells were harvested by centrifugation at 5,000×*g* at 4°C for 15 min, resuspended in lysis buffer (25 mM Tris-HCl, 150 mM NaCl, 20 mM imidazole, pH 9.0) supplemented with 0.1 mg/ml lysozyme, and lysed using a high-pressure homogenizer (LAB 2000; SMT Co., Tokyo, Japan) at 1,000 bar for 10 min. After centrifugation at 10,000×*g* at 4°C for 15 min, the supernatant was loaded onto a Ni-NTA Superflow column (Qiagen, Venlo, Netherlands), and the unbound proteins were washed off with lysis buffer. Subsequently, an overnight culture of BL21 (DE3; pIBA-AtaA_3524-3630_) was diluted 1:100 in LB medium and incubated for 3 h at 37°C. After incubation, anhydrotetracycline (AHTC) (final concentration: 0.2 μg/ml) was added to the medium, and the mixture was further incubated at 28°C for 12 h.

The cells were harvested by centrifugation at 5,000×*g* at 4°C for 15 min, resuspended in lysis buffer supplemented with 0.1 mg/ml lysozyme, and lysed using a high-pressure homogenizer at 1,000 bar for 10 min. After centrifugation at 5,000×*g* at 4°C for 15 min, the supernatant was ultracentrifuged at 100,000×*g* at 4°C for 2 h, and the precipitate was solubilized with 5% Elugent (Merck, Darmstadt, Germany) and suspended in Elugent buffer A (25 mM Tris-HCl, 150 mM NaCl, 20 mM imidazole, 0.5% Elugent, pH 9.0). The suspension was loaded onto a TpgA-N-bound Ni-NTA column, and unbound proteins were washed off with Elugent buffer A. The bound proteins were eluted with Elugent buffer A supplemented with 300 mM imidazole and loaded onto a HiLoad 26/60 Superdex 75 pg column (Cytiva, Marlborough, MA) equilibrated with 25 mM Tris-HCl (pH 7.5) buffer containing 10 mM NaCl and 0.5% Elugent. Peak fractions containing the protein complex were dialyzed using a 12-14 kD MWCO dialysis membrane (Spectrum Laboratories, Rancho Dominguez, CA) against Elugent buffer B (25 mM Tris-HCl, 10 mM NaCl, 0.5% Elugent, pH 9.0) and loaded onto a HiTrap Q HP column (Cytiva) equilibrated with Elugent buffer B. The column was washed three times with 0.6% tetraethylene glycol monooctyl ether (C8E4) containing 25 mM Tris-HCl (pH 9.0) and 10 mM NaCl and two times with 0.6% C8E4 containing 25 mM Tris-HCl (pH 9.0) and 100 mM NaCl. The bound proteins were eluted with 0.6% C8E4 containing 25 mM Tris-HCl (pH 9.0) and 500 mM NaCl, dialyzed against crystallization buffer (25 mM Tris-HCl, 10 mM NaCl, 0.6% C8E4, pH 8.0), and concentrated by ultrafiltration using a 10 kDa cutoff filter device (Amicon Ultra; Merck).

TpgA_24-264_, TpgA_24-138_ (TpgA-N), and TpgA_139-264_ (TpgA-C) were purified as described previously (22). In brief, cells expressing each protein were harvested and lysed using a high-pressure homogenizer. After centrifugation, the protein was purified from the supernatant by Ni-affinity chromatography, ion-exchange chromatography, and size-exclusion chromatography.

### Pull-down assay

For the pull-down assay using recombinant AtaA fragments and cell lysate containing TpgA, an overnight culture of BL21 (DE3) cells harboring plasmids expressing recombinant AtaA fragments was diluted 1:100 in LB medium and incubated for 3 h at 37°C. After incubation, AHTC (final concentration: 0.2 μg/ml) was added to the medium and further incubated at 28°C for 12 h. The cells were harvested by centrifugation at 5,000×*g* at 4°C for 15 min, resuspended in pull-down buffer A (25 mM Tris-HCl, 10 mM NaCl, 0.1% Tween 20, 20 mM imidazole, pH 7.5), and lysed by sonication. After centrifugation at 10,000×*g* at 4°C for 10 min, 500 μl of the supernatant (2 mg/ml) was mixed with 10 μl of Ni-NTA Superflow. After 2 h of incubation on a rotary mixer (NRC-20D; Nissinrika, Tokyo, Japan) at 5 rpm and 4°C, the beads were pelleted by centrifugation at 1,000×*g* for 1 min at 4°C and rinsed three times with pull-down buffer A. The beads were mixed with 500 μl of 5 mg/ml cell lysate from Tol 5 Δ*ataA* and incubated for 2 h on a rotary mixer at 4°C. The beads were pelleted by centrifugation at 1,000×*g* for 1 min at 4°C and rinsed three times with pull-down buffer A. The bound proteins were eluted with 50 μl of 300 mM imidazole (pH 7.5), mixed with 50 μl of 2×SDS‒PAGE sample buffer, and boiled at 98°C for 10 min. The protein samples were separated by SDS‒PAGE and subjected to CBB staining and western blotting analysis with an anti-TpgA antibody (22). For the pull-down assay using recombinant TpgA fragments and cell lysate containing AtaA, 50 μg of His-tagged TpgA_24-264_, His-tagged TpgA_24-138_, or His-tagged TpgA_139-264_ suspended in 300 μl of pull-down buffer A was mixed with 10 μl of Ni-NTA Superflow.

After 1 h of incubation on a rotary mixer at 4°C, the beads were pelleted by centrifugation at 1,000×*g* for 1 min at 4°C, rinsed three times with pull-down buffer A, and resuspended in 300 μl of pull-down buffer A. To prepare the cell lysate of Tol 5 Δ*tpgA*, 100 ml of the cell suspension at an OD_660_ of 0.6 was harvested by centrifugation at 5,000×*g* for 10 min at 4°C and resuspended in 2 ml of BugBuster (Merck), and 50 μg/ml lysozyme and 1 mM phenylmethanesulfonyl fluoride (PMSF) were added. Unlysed cells and debris were removed by centrifugation at 10,000×*g* for 10 min at 4°C. Finally, 200 μl of the cell lysate was mixed with 300 μl of TpgA fragment-bound Ni-NTA beads suspended in pull-down buffer A and incubated on a rotary mixer at 4°C for 2 h. The beads were pelleted by centrifugation at 1,000×*g* for 1 min at 4°C and rinsed with pull-down buffer A for removal of unbound proteins. The bound proteins were eluted with 50 μl of 300 mM imidazole (pH 7.5), mixed with 50 μl of 2×SDS‒PAGE sample buffer, and boiled at 98°C for 10 min. The protein samples were separated via SDS‒PAGE and subjected to CBB staining and western blotting analysis with anti- AtaA_699-1014_ antiserum.

For the pull-down assay using TpgA mutants and cell lysates containing AtaA-TM, 10 μg of His-tagged TpgA variants suspended in 1 ml of pull-down buffer B (25 mM Tris-HCl, 150 mM NaCl, 20 mM imidazole, 0.6% Elugent, pH 7.0) was mixed with 10 μl of Ni-NTA Superflow. After a 1-h incubation on a rotary mixer at 4°C, the beads were pelleted by centrifugation at 1,000×*g* for 1 min at 4°C and rinsed three times with pull-down buffer B. The beads were mixed with 1 ml of 1 mg/ml cell lysate containing AtaA-TM suspended in pull-down buffer B and incubated for 1 h on a rotary mixer at 4°C. The beads were pelleted by centrifugation at 1,000×*g* for 1 min at 4°C and rinsed three times with pull-down buffer B. The bound proteins were then eluted with 25 μl of 500 mM imidazole (pH 7.5), mixed with 25 μl of 2×SDS‒PAGE sample buffer, and boiled at 98°C for 10 min. The protein samples were separated by SDS‒PAGE and subjected to CBB staining.

### Crystallization and X-ray crystallography

The crystallization conditions for the purified protein were first screened by using the sitting-drop vapor-diffusion method at 20°C under 240 conditions using MemPlus (Molecular Dimensions, Sheffield, UK), MemGold (Molecular Dimensions), and The Protein Complex Suite (Qiagen). One microliter of 4.6 mg/ml protein sample suspended in crystallization buffer (25 mM Tris-HCl, 10 mM NaCl, 0.6% C8E4, pH 8.0) and 1 μl of reservoir solution were mixed and periodically observed for 2 weeks under a polarized microscope (BH-2; Olympus). The final optimized crystallization conditions were as follows: 0.2 M MES, 0.4 M calcium chloride, and 35% polyethylene glycol 400 at pH 5.0.

The crystal was loop-mounted and quenched in liquid nitrogen. The diffraction of the crystals was measured under a cryo-condition at the synchrotron beamline NW-12A of the Photon Factory (Tsukuba, Japan) by using a CCD detector (Quantum 270; ADSC, Poway, CA). Diffraction images were processed and scaled by using HKL2000 (40). The CCP4 program suite was used for model building and refinement (41). The phases of all the structures were determined by molecular replacement using MOLREP. The structure was solved using the Hia transmembrane domain (PDB: 2GR7) as the search model. After the molecular replacement, the structures were automatically built by ARP/wARP, manually modeled with Coot, and refined by REFMAC5 and PHENIX (42). The crystallographic parameters are summarized in Table S4. The final coordinates and structure factors that support the findings of this study have been deposited in the PDB with the accession code 9VNJ.

### Bioinformatics

To obtain sequences with similarity to the N-terminal domain of TpgA, the NR70 database was searched for the amino acid sequence of TpgA_24-138_ using PSI-BLAST on the MPI Bioinformatics Toolkit (43). We retrieved 1,461 sequences (E-value <0.001), and the retrieved sequences were clustered using CLANS (32) at a P-value cutoff of 1e-10, an attract value of 10 and a repulse value of 10. The multiple sequence alignment was generated using ClustalO. The DNA sequence of *taa*, which is encoded upstream of the *tpgA*-like gene, was collected from the NCBI nucleotide database. We searched for protein structures similar to that of TpgA-N and calculated the root mean square deviation (RMSD) via the Dali server (44).

### Molecular dynamics (MD) simulations

Using the crystal structure of the protein complex of AtaA-TM and TpgA-N (PDB ID: 9VNJ), two simulation systems were constructed with CHARMM-GUI (45): one consisting of the AtaA-TM and TpgA-N complex, and the other containing only AtaA-TM, with the TM region embedded in a lipid membrane. The lipid membrane was positioned at the center of the simulation box, which was then filled with water molecules. The overall system size was 8.1 nm × 8.1 nm × 16.5 nm. The membrane was asymmetric: the upper leaflet consisted of lipopolysaccharides (LPS) derived from *Acinetobacter baumannii*, with Ca^2+^ ions used to neutralize the negatively charged lipid A and core oligosaccharide regions, while the lower leaflet was composed of a mixture of 1-palmitoyl-2-oleoyl-*sn*- glycero-3-phosphoethanolamine (POPE) and 1-palmitoyl-2-oleoyl-*sn*-glycero-3-phosphatidylglycerol (POPG) in a 4:1 ratio. The LPS in the outer leaflet was modeled as a smooth structure, with the following glycan sequence: α-D-Glc (1→2) β-D-Glc (1→4) β-D-Glc (1→4) β-D-Glc (1→3) [β-D-GalN (1→4)] α-D-GlcNAc (1→4) [β-D-GlcN (1→7)] α-D-Kdo (2→5) [α-D-Kdo (2→4)] α-D-Kdo (2→lipid A). The CHARMM36m force field (46) in combination with the CHARMM-modified TIP3P water model (47) was used to describe all systems. The concentration of sodium chloride ions was set to 0.15 M. All MD simulations were performed using GROMACS 2020 (48). Simulations were conducted under NPT ensemble periodic boundary conditions, where the temperature was controlled at 301.15 K using the V-rescale method (49). The pressure was maintained at 1.0 bar using the Parrinello–Rahman method (50). Bonds involving hydrogen atoms were constrained using the LINCS method (51), and Coulomb interactions were calculated using the particle mesh Ewald (PME) method (52). The Lennard–Jones interactions and real-space PME calculations were set with a cutoff distance of 1.2 nm, and the grid size for the reciprocal-space calculations was set to ensure a grid spacing of 0.1 nm. The time step for all simulations was set to 2 fs. For each system, a 500 ns MD simulation was performed. The RMSD of the atoms corresponding to AtaA-TM was calculated, and the root mean square fluctuation (RMSF) was determined for each residue of AtaA-TM, followed by averaging the RMSF across the three chains.

## Data Availability

Crystallographic coordinates and structure factors have been deposited in the PDB under accession code 9VNJ.

## Acknowledgments

We thank Jens Bassler, Ryuta Hiroshige, Ayumi Hayashi, and Eriko Kawamoto for their technical assistance. We also thank Dirk Linke of the University of Oslo and Masahito Ishikawa of Nagahama Institute of Bio-Science and Technology for helpful discussion. This work was supported by the Japan Society for the Promotion of Science (JSPS) KAKENHI (Grant Numbers JP21H05227 and JP24H00043).

## Author Contributions

K.H. and S.Y. designed the study and wrote the paper. S.Y., A.S., and K.K. prepared the recombinant proteins and conducted the structural analysis. S.Y. and A.L. conducted the bioinformatic analysis. J.S. conducted the molecular simulations. S.Y. and J.K. conducted the other biochemical experiments. All the authors reviewed the manuscript.

