## Supporting Information for "Insights into the complex formation of a trimeric autotransporter adhesin with a peptidoglycan-binding periplasmic protein"

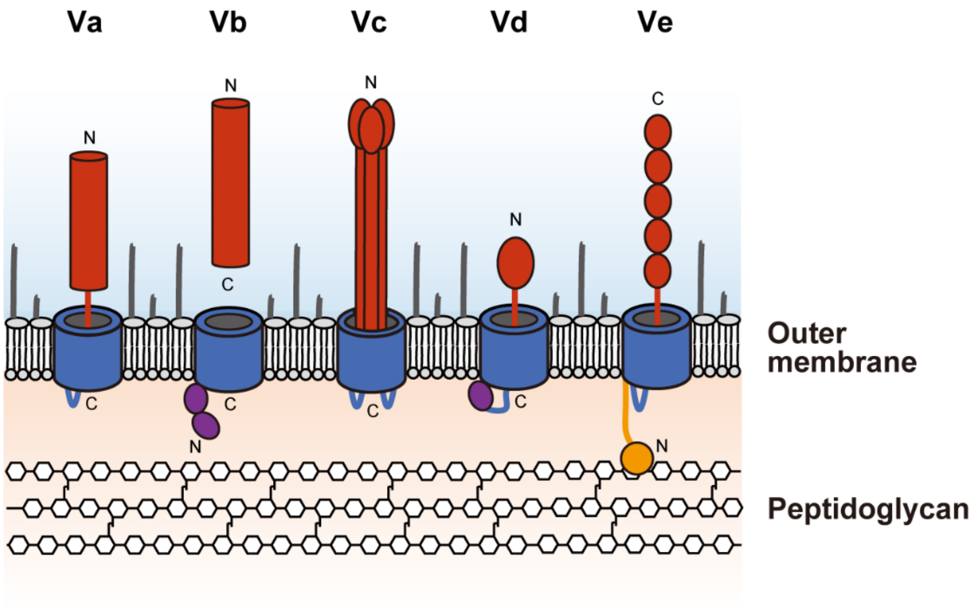


**Figure S1. Schematic representation of type V SSs.** The passenger and transmembrane domains are shown in red and blue, respectively. The POTRA domains and the LysM domain are shown in purple and orange, respectively.

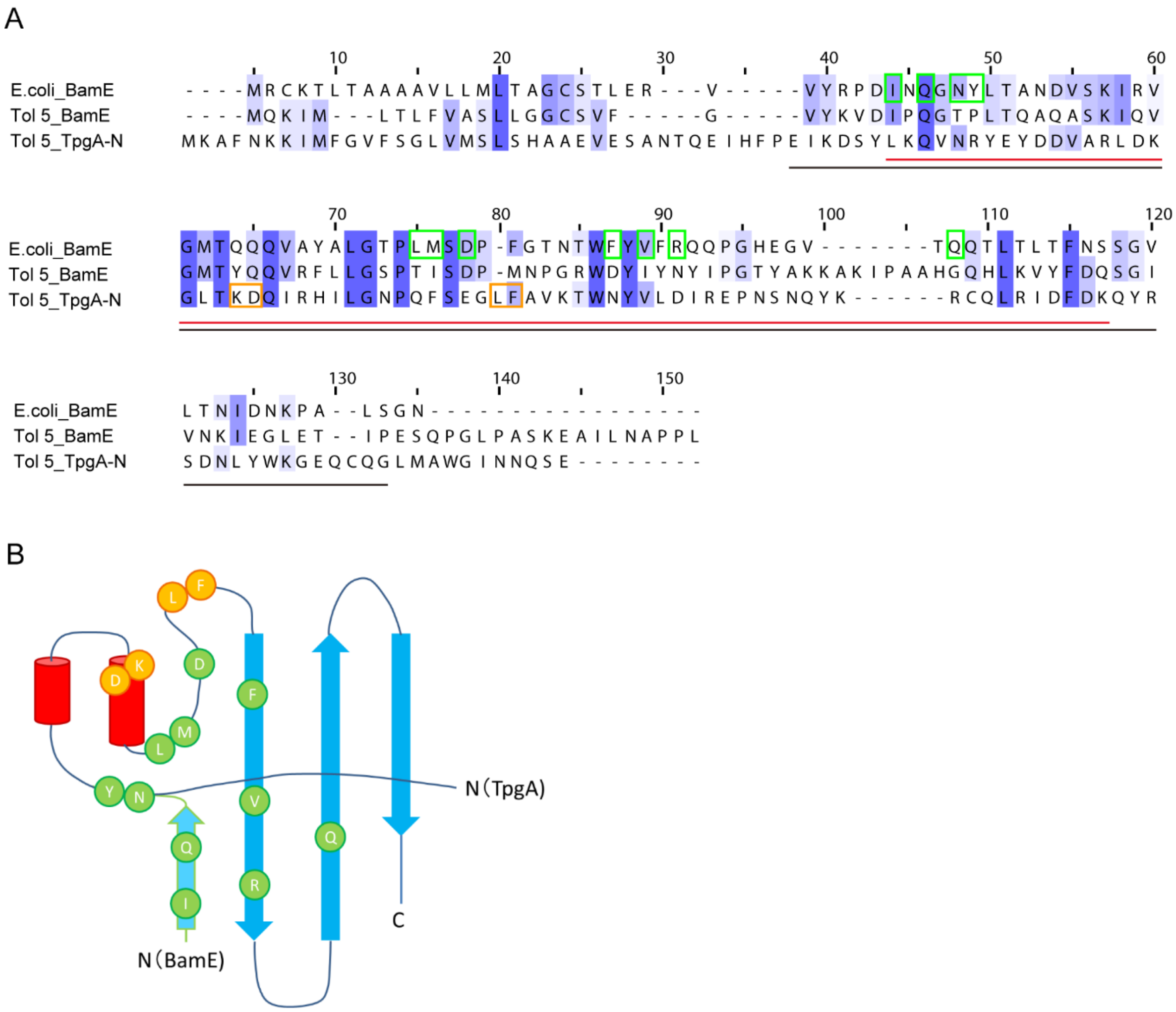


**Figure S2. Comparison of TpgA-N and BamE.** (A) Multiple sequence alignment of BamE from *E. coli*, TpgA-N and BamE from Tol 5. Green and orange boxes indicate residues of *E. coli* BamE that interact with BamA or/and BamD and those of Tol 5 TpgA that interact with AtaA-TM, respectively. The black and red lines indicate the regions used to calculate sequence similarity and structural RMSD. (B) A topology diagram of TpgA-N. α-helices are shown as cylinders, and β-strands are shown as arrows. Interacting residues of *E. coli* BamE with BamA or/and BamD are shown in green and interacting residues of TpgA with AtaA-TM in orange.

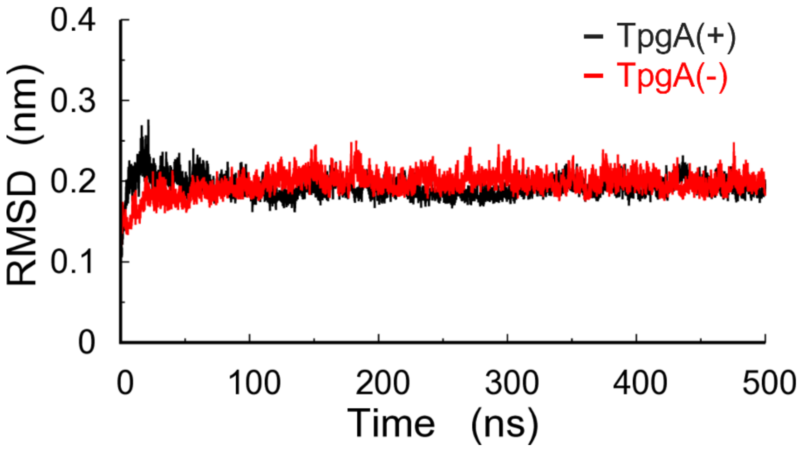


**Figure S3. RMSD of AtaA-TM during MD simulation.**

**Table S1. Genus having *tpgA*-like gene found in cluster I of Figure 4A.**

| Class | Genus | | |
|---|---|---|---|
| Alphaproteobacteria | *Altererythrobacter* | | |
| Betaproteobacteria | *Alcaligenes* | *Achromobacter* | *Advenella* |
| | *Brackiella* | *Burkholderia* | *Collimonas* |
| | *Conchiformibius* | *Eikenella* | *Kerstersia* |
| | *Neisseria* | *Oligella* | *Ottowia* |
| | *Paraburkholderia* | *Ralstonia* | *Variovorax* |
| | *Vitreoscilla* | *Xenophilus* | |
| Gammaproteobacteria | *Acinetobacter* | *Alcanivorax* | *Buttiauxella* |
| | *Chelonobacter* | *Dyella* | *Enterobacter* |
| | *Erwinia* | *Escherichia* | *Frateuria* |
| | *Halomonas* | *Klebsiella* | *Luteimonas* |
| | *Lysobacter* | *Moraxella* | *Oblitimonas* |
| | *Oleiagrimonas* | *Pasteurella* | *Psychrobacter* |
| | *Pseudomonas* | *Pseudoxanthomonas* | *Rhodanobacter* |
| | *Salmonella* | *Serratia* | *Stenotrophomonas* |
| | *Xanthomonas* | *Yokenella* | |
| Campylobacteria | *Campylobacter* | | |

**Table S2. Bacterial strains and plasmids used in this study.**

| Strain | Description | Reference |
|---|---|---|
| *Acinetobacter* sp. Tol 5 | Wild type strain, expressing *ataA* | (1) |
| *Acinetobacter* sp. Tol 5 Δ*ataA* | *Acinetobacter* sp. Tol 5 4140, Unmarked Δ*ataA* mutant of Tol 5, *ataA*$^-$ | (2) |
| *Acinetobacter* sp. Tol 5 Δ*tpgA* | *Acinetobacter* sp. Tol 5 OM186, Unmarked Δ*tpgA* mutant of Tol 5, *tpgA*$^-$ | (3) |
| *Escherichia coli* DH5α | Host for cloning | Takara Bio |
| *Escherichia coli* BL21(DE3) | Host for expression of recombinant proteins | Thermo Fisher Scientific |
| *Escherichia coli* S17-1 | Donor strain for bacterial conjugation | (4) |
| Plasmids | | |
| pIBA-GCN4tri-His | Expression vector for recombinant AtaA proteins, Amp$^r$ | (5) |
| pASK-IBA12 | Expression vector for secreted proteins, Amp$^r$ | IBA Lifesciences |
| pET28d | Expression vector for His-tagged protein, Km$^r$ | (3) |
| pARP3 | *E. coli-Acinetobacter* shuttle expression vector, Gm$^r$, Amp$^r$ | (6) |
| pAtaA | *ataA*-expression vector, pARP3::*ataA* | (6) |
| pTpgA | *tpgA*-expression vector, pARP3::*tpgA* | (3) |

**Table S3. Primers used in this study.**

| Name | Sequence (5'-3') |
|---|---|
| AtaA723-f | TGCAAGGGTCTCCGATTGACTCAAAAGCAGCAGCATC |
| AtaA806-r | TGCAGCGGTCTCCTCATAGCATTTTTGCATCTACAC |
| AtaA1221-f | TGCAAGGGTCTCCGATTGATGATGCAGGTACAGCATTAAC |
| AtaA1416-r | TGCAGCGGTCTCCTCATGTCACCTGCGTTTACACCTT |
| AtaA2215-f | TGCAAGGGTCTCCGATTAACAATGCAGTTGTTGATG |
| AtaA2395-r | TGCAGCGGTCTCCTCATTTGGCTACCATTGATCGC |
| AtaA2777-f | TGCAAGGGTCTCCGATTTACGATATTGCAAACAGCG |
| AtaA2903-r | TGCAGCGGTCTCCTCATTAACTGTTCTTCAGTCGCAG |
| AtaA3524-f | AGGCCGCTAGCCATCATCATCATCATCATGCAGATCAAGCATTGAA TAGC |
| AtaA3630-r | GGTATCAGCGGTGTGATTGACTAAAAGCTTGACCTGTG |
| TM-dHis-f | GCAGATCAAGCATTGAATAGC |
| TM-dHis-r | GCTAGCGGCCTGCGCTAC |
| TpgA-N-f | AGTCTGAGGCGGCCGCACTCGAGCAC |
| TpgA-N-r | GCGGCCGCCTCAGACTGATTATTAATCCCC |
| TpgA-C-f | GATATACCATGACTGAGCAAACGACTCTAG |
| TpgA-C-r | GTTTGCTCAGTCATGGTATATCTCCTTCTTAAAC |
| TpgA-64-f | ACCGCAGATCAGATTCGCCATATTTTGG |
| TpgA-64-r | AATCTGATCTGCGGTTAATCCCTTGTCTAAAC |
| TpgA-65-f | ACCAAAGCACAGATTCGCCATATTTTGG |
| TpgA-65-r | AATCTGTGCTTTGGTTAATCCCTTGTCTAAAC |
| TpgA-64/65-f | ACCGCAGCACAGATTCGCCATATTTTGG |
| TpgA-64/65-r | AATCTGTGCTGCGGTTAATCCCTTGTCTAAAC |
| TpgA-80-f | GAAGGTGCATTTGCGGTTAAGACATGG |
| TpgA-80-r | CGCAAATGCACCTTCAGAGAATTGAGGATTTC |
| TpgA-81-f | GGTCTTGCAGCGGTTAAGACATGGAATTATG |
| TpgA-81-r | AACCGCTGCAAGACCTTCAGAGAATTGAGG |
| TpgA-80/81-f | GGTGCAGCAGCGGTTAAGACATGGAATTATG |
| TpgA-80/81-r | AACCGCTGCTGCACCTTCAGAGAATTGAGGATTTC |

**Table S4. Data collection and refinement statistics of the N-terminal domain of TpgA and the transmembrane domain of AtaA.**

| | |
|---|---|
| PDB code | 9VNJ |
| Space group | R32 |
| a, b, c (Å) | 61.94, 61.94, 428.9 |
| α, β, γ (º) | 90, 90, 120 |
| Resolution range (Å) | 37.92-2.60 (2.64-2.60) |
| Wavelength (Å) | 1.0 |
| Completeness (%) | 98.45 (99.0) |
| Redundancy | 8.4 (7.9) |
| $CC_{1/2}$ | 0.971 |
| $I/\sigma(I)$ | 45.6 (9.3) |
| $R_{merge}$ (%) | 7.5 (35.5) |
| Unique reflections | 10,168 (477) |
| $R_{work}/R_{free}$ (%) | 19.5/24.4 |
| Total number of atoms | 1788 |
| Wilson B-factor (Å$^2$) | 40.1 |
| Ramachandran statistics (%) (preferred/allowed/outliers) | 97.0/2.5/0.5 |
| Bond length/angle RMSD (Å/º) | 0.013/1.169 |
| Average B, all atoms (Å$^2$) | 42.0 |

* Values in parenthesis refer to the highest-resolution shell (2.64-2.60 Å).
